# Epistasis reduces fitness costs of influenza A virus escape from stem-binding antibodies

**DOI:** 10.1101/2022.07.14.500125

**Authors:** Chung-Young Lee, C. Joaquin Caceres, Ginger Geiger, Brittany Seibert, Flavio Cargnin Faccin, L. Claire Gay, Lucas M. Ferreri, Drishti Kaul, Jens Wrammert, Gene S. Tan, Daniel R. Perez, Anice C. Lowen

## Abstract

The hemagglutinin (HA) stem region is a major target of universal influenza vaccine efforts owing to the presence of highly conserved epitopes across multiple influenza A virus strains and subtypes. To explore the potential impact of vaccine-induced immunity targeting the HA stem, we examined the fitness effects of viral escape from stem-binding broadly neutralizing antibodies (stem-bnAbs). Recombinant viruses containing each individual antibody escape substitution showed diminished replication compared to wild-type virus, indicating that stem-bnAb escape incurred fitness costs. A second-site mutation in the HA head domain (N133D) reduced the fitness effects observed in primary cell cultures and likely enabled the selection of escape mutations. This putative permissive mutation was not, however, sufficient to ease fitness costs in a ferret transmission model. Taken together, these data suggest that viral escape from stem-bnAbs is costly but highlight the potential for epistatic interactions to enable evolution within the functionally constrained HA stem domain.

## Introduction

Hemagglutinin (HA) is the surface glycoprotein of influenza A virus (IAV) that mediates viral attachment and entry into host cells. The globular head of HA contains the receptor binding site (RBS) that binds to sialylated glycans on respiratory epithelial cells, allowing their uptake by endocytosis^1, 2^. During the endocytic pathway, an acidic endosomal environment triggers release of the fusion peptide from within the stem domain of the HA trimer, leading to fusion between viral and endosomal membranes and release of the viral ribonucleoproteins (vRNPs) into the cytoplasm^3, 4, 5^.

The globular head of HA is the main target of neutralizing antibodies elicited by vaccination or infection^6, 7^. These antibodies typically act by blocking the interaction between HA and sialylated glycan receptors, inhibiting viral attachment^5, 8, 9^. However, selective pressure leads to antigenic changes in circulating viruses, causing reduction in or loss of neutralization^10, 11, 12^. This high sequence variability in HA lowers the cross-protection of humoral immune responses elicited by traditional vaccines^13^. The resultant narrow specificity of influenza vaccines necessitates their annual updating and gives the potential for mismatch between circulating strains and those included in the vaccine. Therefore, substantial efforts have been invested to develop vaccines targeting more conserved regions of IAV, with the goal of conferring broad protection^7^.

The stem region in HA has considerably higher sequence conservation than the globular head. The stem consists of the N-and C-terminal regions of HA1 and most of HA2. Generally, antibodies to the stem are harder to elicit but have greater breadth than head binding antibodies, neutralizing a broader range of IAVs^6, 14, 15, 16, 17^. In addition, the stem region is thought to be inherently less permissive to viral escape because it is essential to stabilize the HA trimeric structure and to drive viral fusion through HA conformational change^18, 19^. Therefore, the stem has been regarded as a promising target for universal vaccines and anti-viral therapeutics ^7^. Several stem-targeted, broadly reactive, neutralizing antibodies (stem-bnAbs) have been isolated from infected or immunized individuals since the first stem-bnAb was isolated in 1993^14^. These antibodies are categorized based on the breadth of their neutralization potential to group 1 IAVs^14, 15, 20, 21, 22^, group 2 IAVs^17, 23^, pan-IAV^16, 24^, or IAVs and IBVs^25^.

High conservation of the HA stem domain within circulating viruses suggests that its evolution is limited due to functional or structural constraints. Laboratory-based studies in which selection on the stem domain is imposed experimentally can give valuable insight. In this context, a few escape mutations against stem-bnAbs have been identified in different subtypes of IAV ^15, 26, 27, 28^. However, most conferred a fitness cost, attenuating viral replication in either *in vitro* or *in vivo* models ^26, 28, 29^. This attenuation is likely associated with lower mutational tolerance of the HA stem domain compared to the HA head domain^30^, even under immunological pressure^31^.

In this study, we monitored the evolution of influenza A/Netherlands/602/2009 (NL09) virus, a 2009 pandemic H1N1 strain, in the presence of two different stem-bnAbs (70-1F02 and 05-2G02). We identified several individual escape mutations in this process. As expected, all isolated escape mutations carried viral fitness costs. However, these costs were diminished by a second-site mutation, N133D, which was found during virus passage prior to fixation of the escape mutations. N133D increased virus binding to host cell receptors, resulting in enhanced replication of the escape mutant viruses in primary human cell cultures. Conversely, this putative permissive mutation did not enable improved replication and transmission of the mAb-insensitive strains in ferrets, suggesting that the evolution of stem variant viruses may be more constrained *in vivo* than in cell culture models. These data emphasize the importance of positive epistasis for viral evolution under selective pressure but overall support the argument that HA stem-targeted vaccines would be relatively robust to antigenic drift.

## Results

### Identification of mutations allowing escape from two stem-bnAbs

To isolate escape variants against stem-bnAbs, A/Netherlands/602/2009 (H1N1) (NL09) virus was serially passaged ten times with either 70-1F02 or 05-2G02 stem-bnAbs as described before^32^. 70-1F02, derived from an individual naturally infected with a pandemic H1N1 IAV, has demonstrated cross-reactivity to H1 and H5 subtype strains^20, 33^. 05-2G02, isolated from a healthy subject immunized with the inactivated 2009 pandemic H1N1 vaccine, has broad reactivity with strains of group 1 and group 2 HAs^24, 33^. As a control, we also passaged NL09 virus without stem-bnAbs ten times (Supplementary Fig. 1). Each passage regimen was carried out in triplicate.

To assess the sensitivity of the passaged virus populations to stem-bnAbs, 50% plaque reduction neutralizing titers (PRNT_50_) were measured (Fig. 1). The PRNT_50_ titers of viral populations passaged with stem-bnAbs gradually increased with passage number and all six passage-ten populations showed titers of more than 100 μg/mL.

**Fig. 1.**
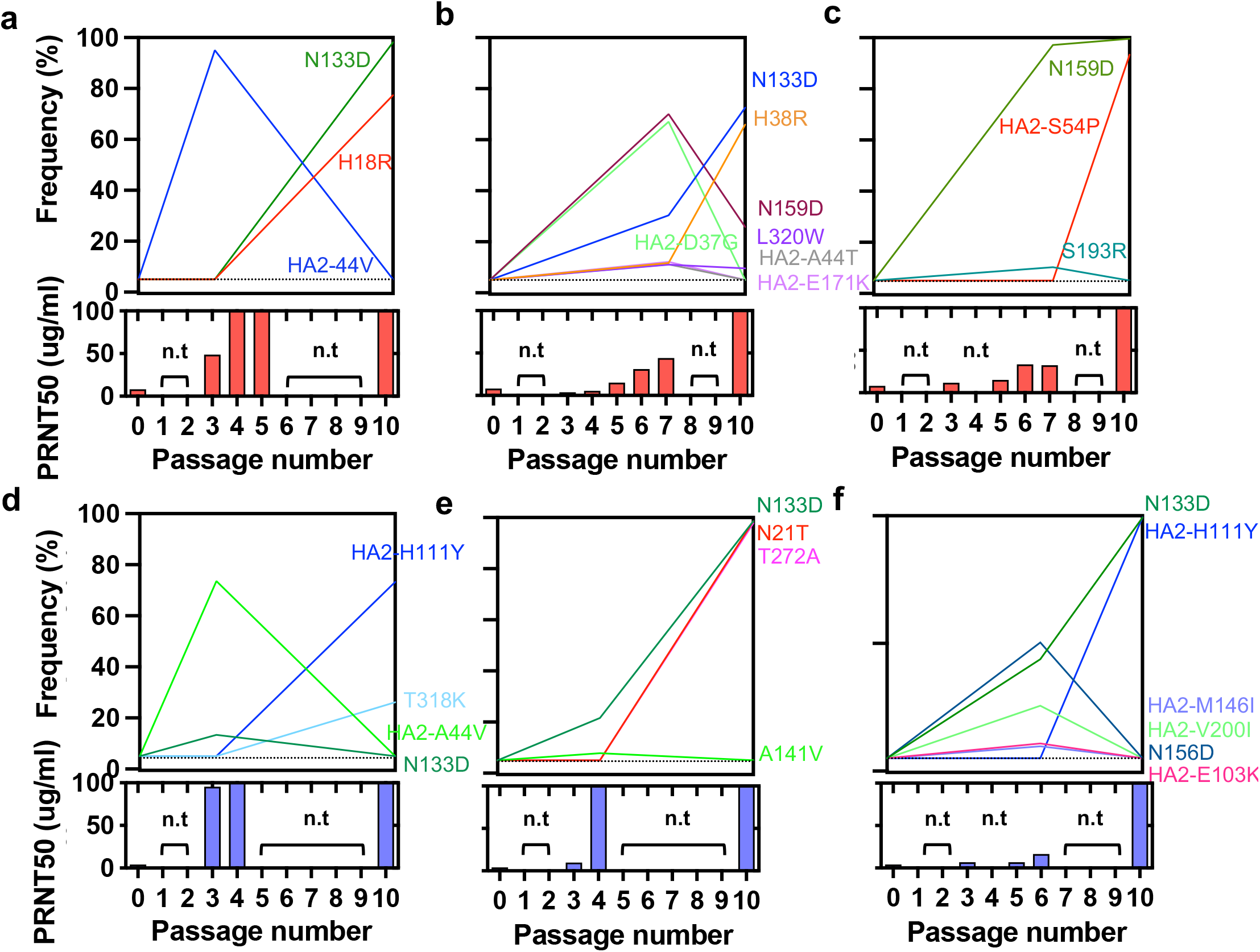
Dynamics of mutation frequency and resistance of passaged NL09 virus populations to stem-bnAbs. NL09 virus was combined with (a-c) 70-1F02 or (d-f) 05-2G02 antibody and serially passaged 10 times with increasing concentrations of antibody. A single replicate was performed with an initial concentration of 20 (a and d), 2 (b and e), or 1 (c and f) μg/mL of 70-1F02 or 05-2G02, for a total of three replicates per stem-bnAb. n.t = not tested; dotted line = limit of detection (5%). The variants that showed PRNT_50_ titers >30 μg/mL were sequenced.

To identify and monitor the dynamics of escape mutations, deep sequencing was conducted on virus populations derived from both early and late passages. In addition, to confirm the frequency of mutations and assess their co-occurrence, we sequenced six plaque-purified clones per sample. The results revealed that, within each passaged lineage, a single variant harboring one to three mutations became dominant. Each variant contained distinct mutations although some were shared (Fig. 1). The substitution N133D (H3 numbering)^34^ was dominant in four out of six antibody selected populations and one out of four controls. H18R and H38R in HA1 and S54P in HA2 became dominant in the populations passaged with 70-1F02. By contrast, N21T and T272A in HA1 and H111Y in HA2 became dominant in the populations passaged with 05-2G02. To identify single mutations that confer antibody escape, we generated recombinant NL09 viruses bearing the individual mutations detected in the passaged virus variants, and measured viral resistance to antibody (Fig. 2a,b). The recombinant viruses harboring H18R, H38R, L320W, or HA2-S54P are highly resistant to 70-1F02 antibody, showing PRNT_50_ titers of more than 300 μg/mL. Similarly, the mutant viruses harboring T318K or HA2-H111Y are highly resistant to 05-2G02 antibody, showing titers of >300 or 202.9 μg/mL, respectively. Of note, the mutations conferring high PRNT_50_ titers coincide with those that dominated in passage ten populations, suggesting that their phenotypic effects led to their positive selection.

**Fig. 2.**
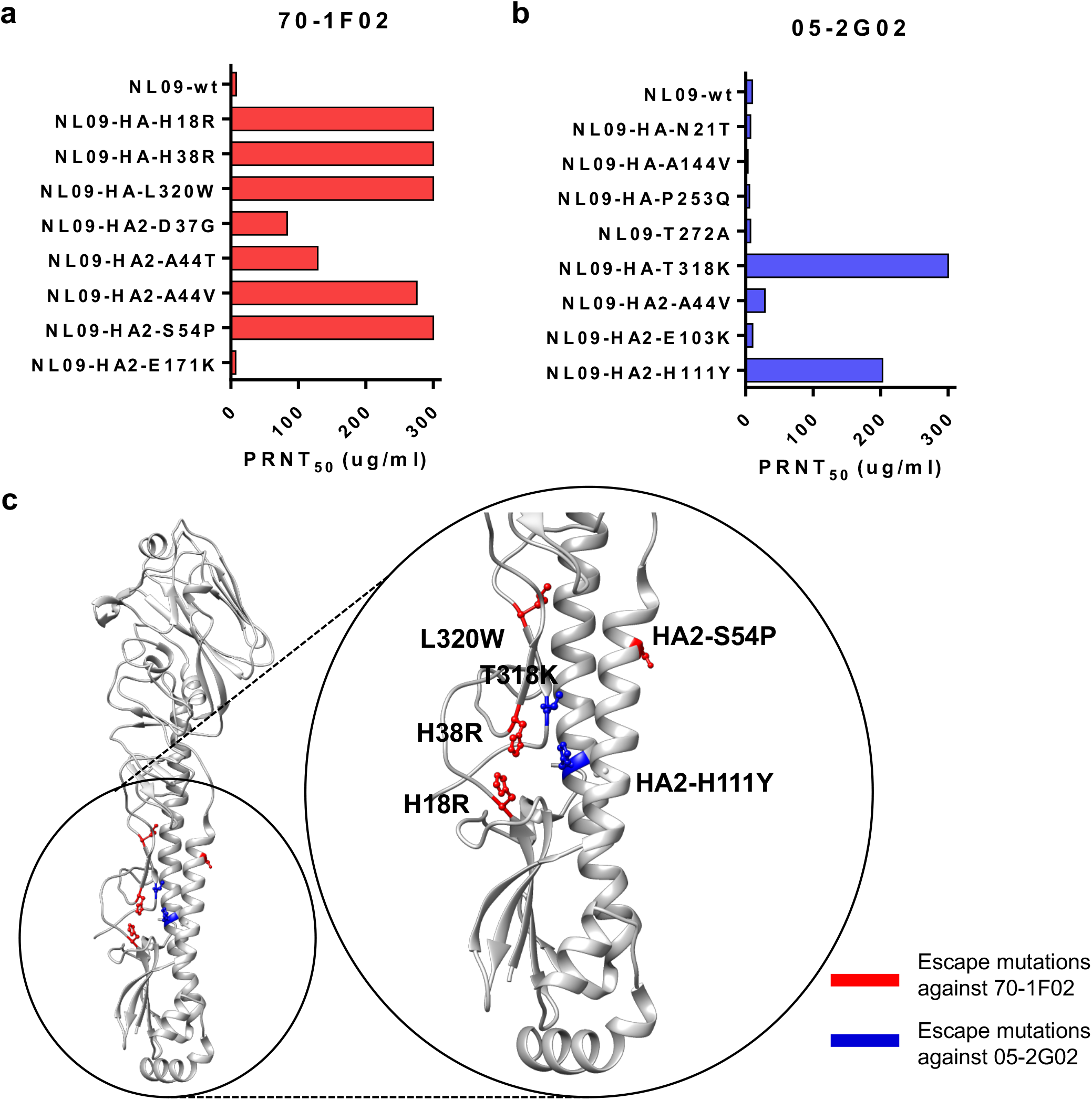
Identification of stem-bnAb escape mutations. (a and b) PRNT_50_ titers of recombinant NL09 viruses harboring individual mutations found in the passaged NL09 variants. Concentrations of (a) 70-1F02 and (b) 05-2G02 antibodies leading to a 50% reduction in plaque counts are shown. (c) Locations of selected escape mutations on the HA of A/California/04/2009 (H1N1) (PDB ID 3LZG)^48^.

### The escape mutations decrease antibody binding to NL09 HA

To investigate the mechanisms by which the escape mutations in HA confer resistance, we first mapped them onto a pandemic H1N1 HA structure (PDB ID 3LZG). The mutations conferring high resistance to the stem-bnAbs are located in HA stem region (Fig. 2c). Among these mutations, L320W and HA2-S54P are buried within the monomer interior, and H18 and H38 are present within the 70-1F02 epitope^33^. Because a common strategy of escape is to abolish or reduce binding between antigen and antibody, we next evaluated the impact of the escape mutations on antibody binding. An ELISA-based assay was used in which NL09 HA proteins harboring an individual escape mutation were over-expressed in adherent cells. HA protein expression was confirmed quantitatively using an HA head specific antibody (1009-3E04). Both 70-1F02 and 05-2G02 stem-bnAbs bound to the cells expressing NL09 HA-wt (Fig. 3a,b). However, binding to 70-1F02 was completely abolished by H18R, H38R, and L320W mutations and significantly reduce by HA2-S54P mutation. Similarly, binding to 05-2G02 was eliminated by T318K and HA2-H111Y mutations.

**Fig. 3.**
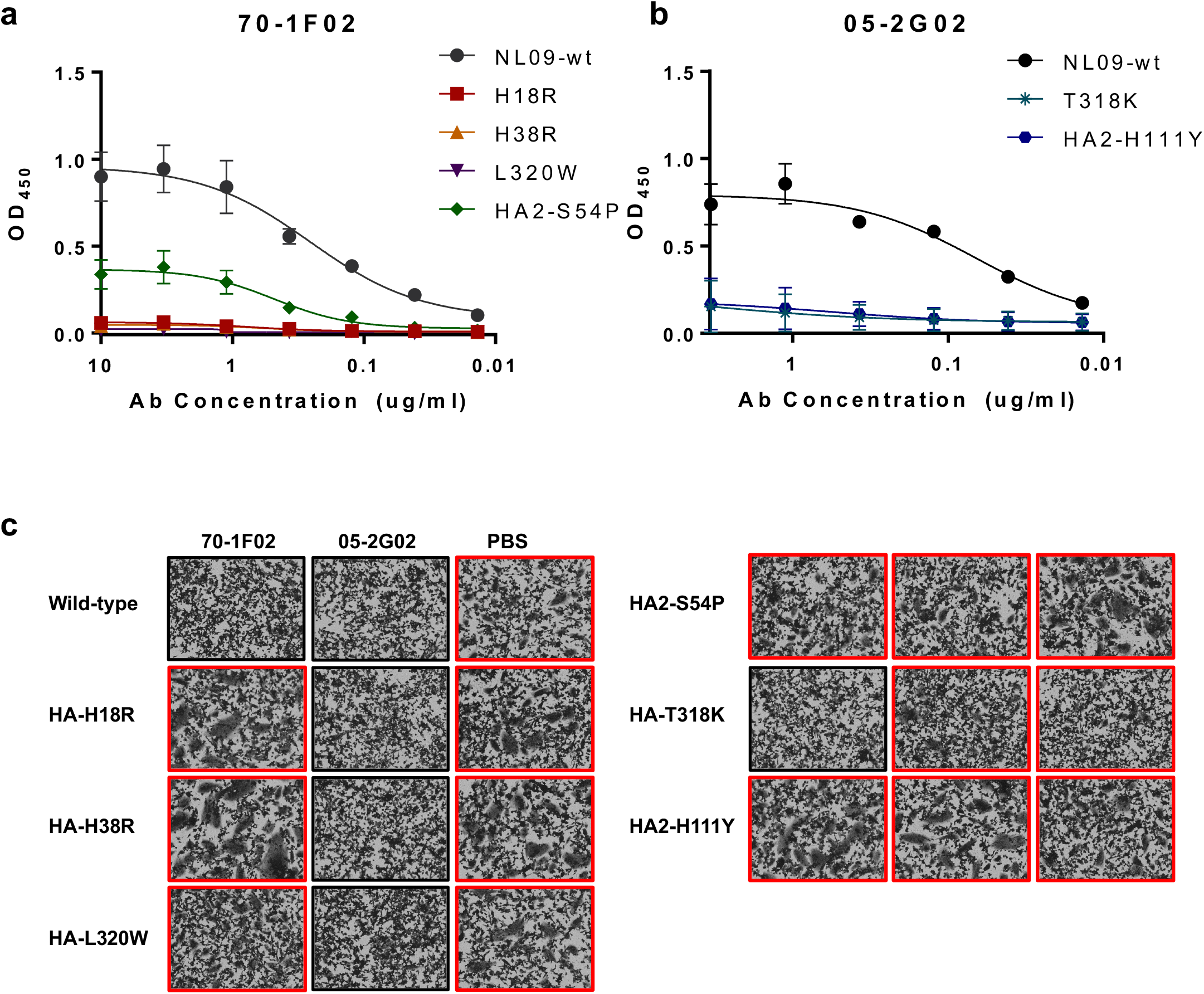
By abolishing antibody binding, the escape mutations facilitate HA-membrane fusion in the presence of stem-bnAbs. (a and b) Binding between mutant HA and the stem-bnAbs (a) 70-1F02 or (b) 05-2G02. (c) HA mediated membrane fusion was evaluated by monitoring for syncytia formation at low pH. Red rectangles indicate that syncytia formation was observed.

Next, we investigated whether the loss of interaction between HA and antibody affects viral fusion activity using a syncytia formation assay. In NL09 HA-wt, both stem-bnAbs inhibited fusion (Fig. 3c). Conversely, cells over-expressing mutant HA proteins demonstrated syncytia formation in the presence of stem-bnAb. Notably, the HA2-S54P and H111Y mutants formed syncytia in the presence of both 07-1F02 and 05-2G02, suggesting that these mutations confer resistance to both 70-1F02 and 05-2G02, although it was selected by either 70-1F02 or 05-2G02. These data indicate that each escape mutation examined acts by interrupting the binding of stem-bnAbs, enabling functional HA-membrane fusion in the presence of antibody.

### The escape mutations confer fitness defects

Given high conservation of the HA stem domain, mutations in the stem were expected to carry fitness costs. To test for fitness effects, we first measured the growth kinetics of NL09 viruses harboring the individual escape mutations in MDCK, A549, and NHBE cells. In MDCK cells, the NL09 recombinant viruses carrying the individual escape mutations replicated comparably to NL09-wt virus, although viruses harboring T318K, L320W, HA2-S54P, or HA2-H111Y were slightly attenuated (Fig. 4a). Notably, stronger phenotypes were observed in A549 and NHBE cells (Fig. 4b,c). In A549 cells, the viruses harboring T318K, L320W, and HA2-S54P replicated poorly compared to NL09-wt virus, while replication of NL09-HA-H18R, NL09-HA-H38R, NL09-HA2-H111Y viruses was comparable to NL09-wt. In NHBE cells, all six mutant viruses were significantly attenuated, although H18R, H38R, and HA2-H111Y were more fit than the other mutant viruses. Collectively, the impact of the single escape mutations on viral fitness varied widely and was most readily detected in NHBE cells.

**Fig. 4.**
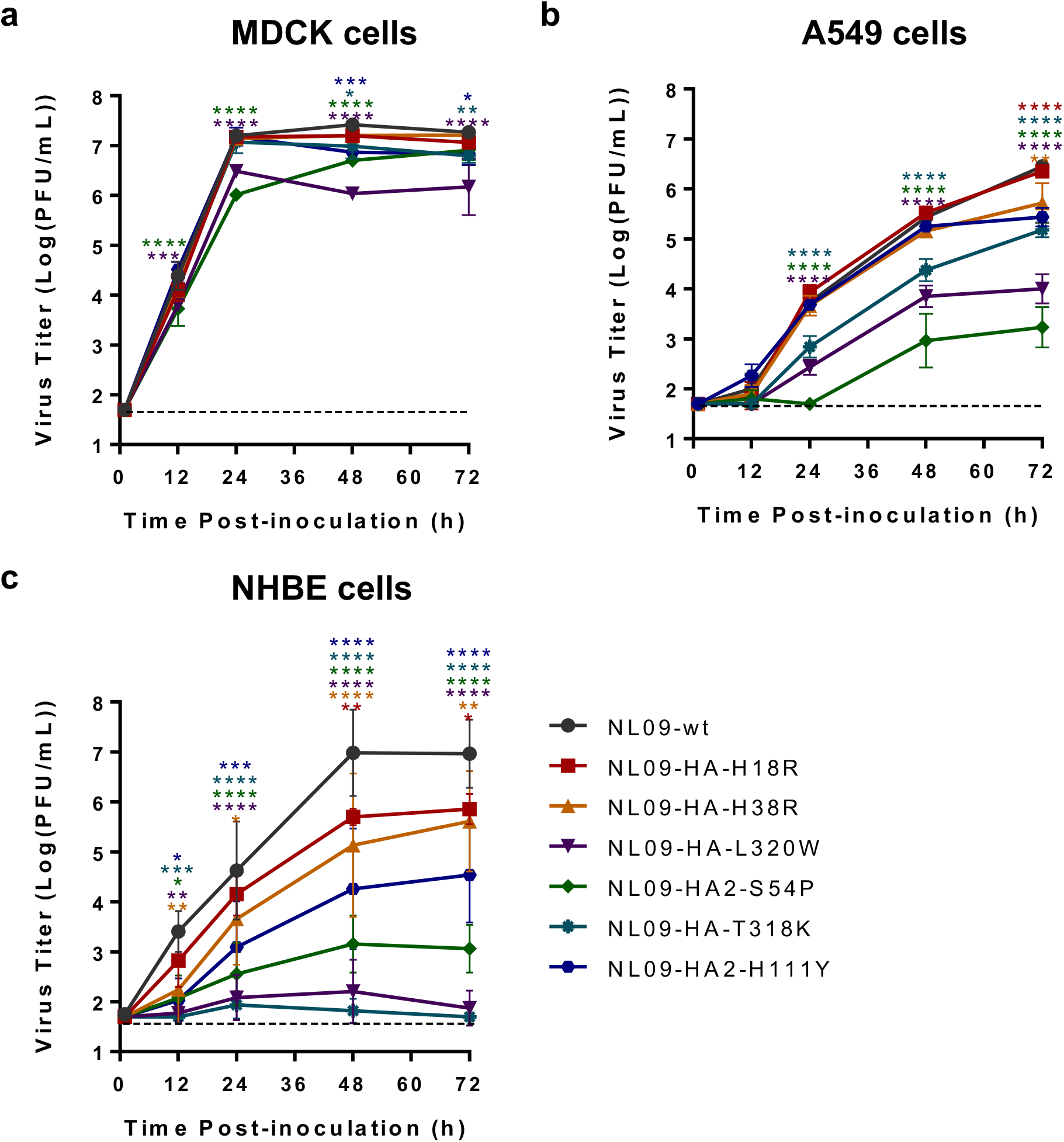
Growth kinetics of recombinant viruses harboring individual escape mutations. (a) MDCK cells were infected at an MOI of 0.002 PFU/cell; (b) A549 cells and (c) NHBE cells were infected with MOI=0.01 PFU/cell. Mean results ± s.d. of at least three independent experiments are plotted. Statistical significance was assessed by two-way ANOVA with Sidak’s multiple comparisons test. ****, < 0.0001; ***, < 0.001; **, < 0.01; *, < 0.05.

These findings prompted us to examine mechanism by which the escape mutations impact viral fitness. Owing to their location in the HA stem and the stem’s role in HA-mediated membrane fusion, we assessed pH of fusion and fusion kinetics. First, a syncytia formation assay was employed to measure the pH at which fusion takes place (Supplementary Fig. 2a). The HAs harboring the individual escape mutations led to membrane fusion at pH 5.3 or 5.4, a minor difference from NL09-wt HA. Next, fusion kinetics of the mutant HAs were evaluated. For this purpose, a head specific antibody that is completely detached by acid-induced HA conformational change (Supplementary Fig. 3) was used to monitor HA conformation over a two minute time course. The half-life for antibody detachment from NL09-wt HA was 4.6 s (Supplementary Fig. 2b,c). T318K, L320W and HA2-H54P mutations decreased the half-life (to 2.9, 3.9, and 2.4 s, respectively), indicating that these mutations destabilize the pre-fusion HA. This observation is consistent with the more severe attenuation of replication seen with these mutations. Although HA2-H111Y mutation (half-life = 2.9 s) showed faster fusion kinetics then NL09-wt, H18R (half-life = 4.2 s) and H38R (half-life = 5.0 s) mutations exhibited similar fusion kinetics to NL09-wt. The modest effects on fusion seen for these three mutants is again consistent with their intermediate attenuation in cell culture. Taken together, the escape mutations identified have minimal impact on the pH of fusion but those with strong fitness effects hastened the fusion event, accounting for the attenuated replication observed.

Since the escape mutations were selected under antibody treatment, we evaluated their fitness relative to NL09wt in the presence of stem bnAbs. NHBE cells were pre-treated with 1 PRNT_50_ of the stem-bnAbs or PBS, then inoculated with MOI=1 of NL09-wt or the single escape mutant viruses. As expected, the replication of NL09-wt virus significantly decreased in the presence of 07-1F02 or 05-2G02 antibodies (Supplementary Fig. 4). In contrast, the escape mutant viruses harboring H18R, H38R, or HA2-H111Y replicated similarly with or without the antibodies, confirming their resistance to the relevant immunological pressure.

### N133D, a potential permissive mutation, restores viral fitness in human cell cultures

In addition to the escape mutations, four out of six passage 10 escape variants have a common second site mutation (N133D, H3 numbering) in the HA globular head domain. Residue 133 is located in 130-loop, between Ca2 and Sa antigenic sites (Supplementary Fig. 5a). It is close to the receptor binding site and has been reported to affect antigenicity and receptor binding affinity^35, 36^. Of note, since 2019, virus strains harboring an N133D mutation have increased in frequency and become dominant within the circulating 2009 pandemic H1N1 lineage (Supplementary Fig. 5b). Given these prior observations, we hypothesized that N133D may act as a permissive mutation and tested its potential to modulate the fitness of the escape mutant viruses.

First, we evaluated the impact of HA N133D on sensitivity to stem-bnAbs and HA fusion potential. The recombinant virus carrying N133D showed similar PRNT_50_ titer, fusion pH, and fusion kinetics compared to NL09-wt (Supplementary Fig. 5).

Next, we examined the impact of HA N133D on receptor preference by evaluating binding of NL09-wt and NL09-HA-N133D viruses on a glycan array (Supplementary Fig. 5a). Both viruses bound predominantly to Sia-α2,6 glycans. In addition, extensive overlap in bound glycans was seen, with seven of the top ten glycans common between NL09-wt and NL09-HA-N133D viruses (Fig. 5a). The data suggest that N133D makes little change to the HA receptor binding profile.

**Fig. 5.**
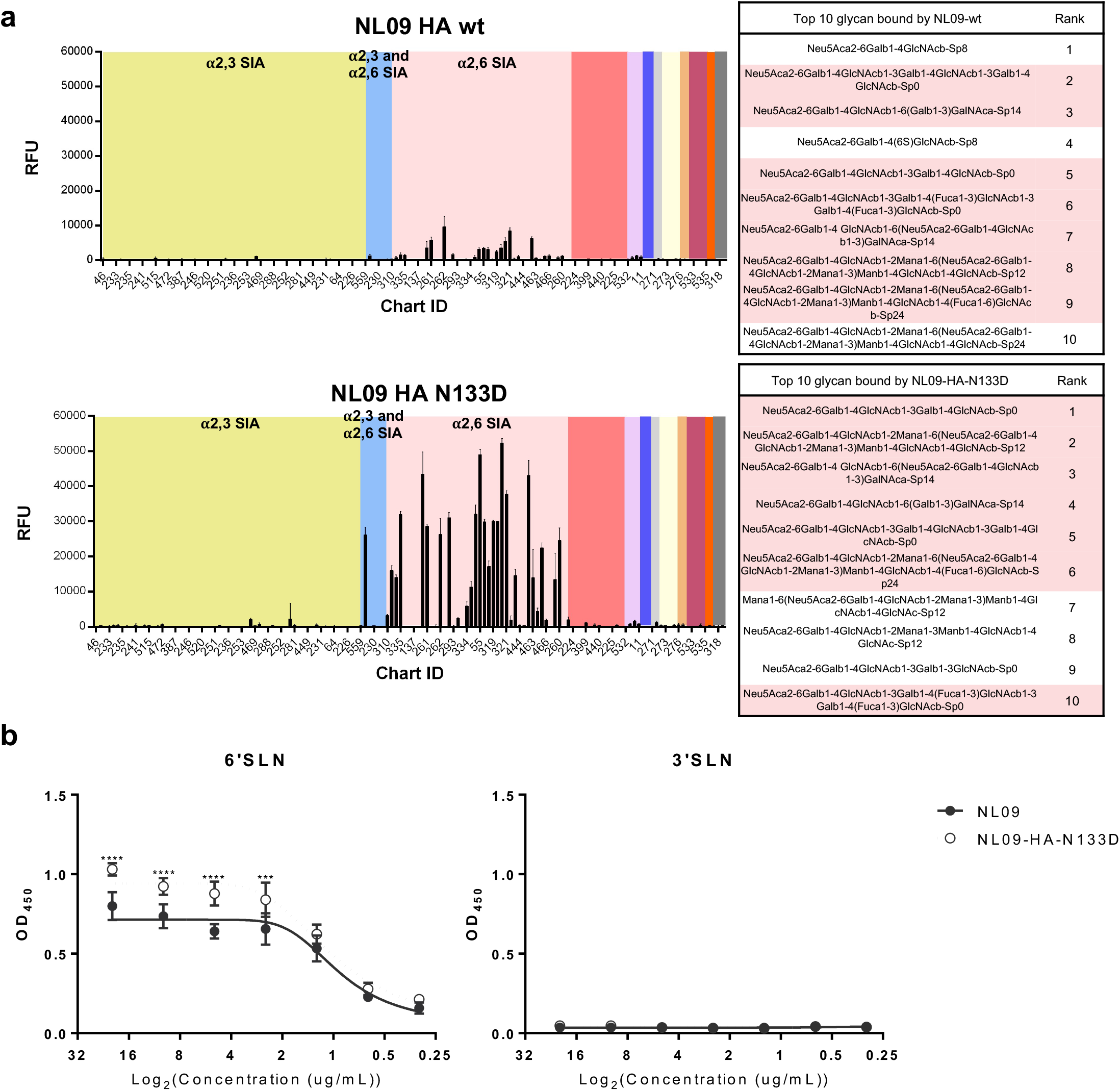
N133D mutation in HA does not affect viral receptor binding preferences but increases strength of receptor binding. (a) Profiles of virus binding to v4.0 of the CFG glycan array are shown as the relative intensity of the fluorescence (RFU) for each glycan structure, indicating virus recognition and binding. The different linkage of sialylated glycans are highlighted by yellow=2,3 SIA, blue=2,3 and 2,6 SIA, and pink=2,6 SIA. (b) The binding of each virus to human-like (6’SLN) and avian-like (3’SLN) were measured by solid-phase binding assay. The data shown were the mean ± s.d. with at least three independent experiments. Statistic significance was analyzed by two-way ANOVA with Sidak’s multiple comparisons test. ****, < 0.0001; ***, < 0.001.

We then tested whether the N133D mutation modulates the strength of IAV receptor binding using solid-phase binding assay (Fig. 5b). Attachment of NL09-wt virus to an avian-like receptor (3’SLN) was at the limit of detection and the addition of N133D did not enhance this binding. In contrast, NL09-wt virus bound robustly to a human-like receptor (6’SLN), as expected. Notably, compared to NL09-wt virus, NL09-HA-N133D virus showed higher levels of binding to this human-like receptor. These data suggest that the addition of the N133D mutation to HA does not affect the receptor binding profile but enhances binding to a human-like receptor.

Examination of the impact of HA N133D on viral replication revealed that the NL09-HA-N133D virus achieved higher titers than NL09-wt virus in NHBE cells (Fig. 6a). To elucidate the effects of N133D on viral fitness in the context of escape mutations in the stem domain, recombinant viruses containing N133D and one of three individual escape mutations (NL09-HA-H18R-N133D, NL09-HA-H38R-N133D, and NL09-HA2-H111Y-N133D) was evaluated (Fig. 6). Similar to NL09-HA-N133D virus, higher viral replication was observed in the viruses carrying the dual mutations compared to the viruses carrying the single mutation in NHBE cells. Notably, the growth of the viruses carrying dual mutations was comparable to that of NL09-wt virus, suggesting that co-existence of the second-site mutation with the escape mutation reduces viral fitness costs.

**Fig. 6.**
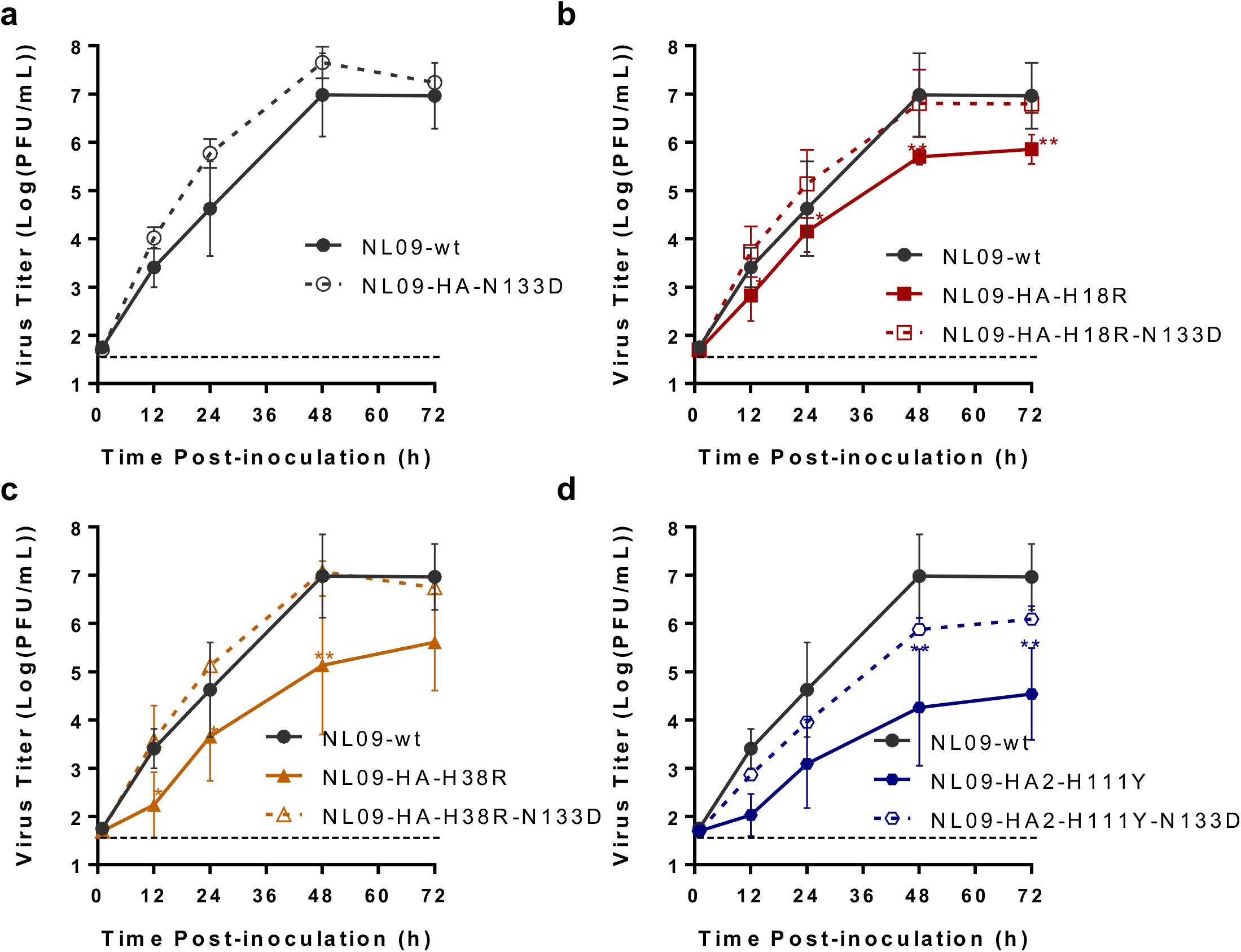
Growth kinetics of recombinant NL09 viruses harboring individual escape mutation and N133 D in NHBE cells. (a-d) Data plotted are the mean ± s.d of at least three independent experiments. Statistical significance was assessed using a two-way ANOVA with Sidak’s multiple comparisons test. **, < 0.01; * < 0.05.

### N133D is not sufficient to restore viral fitness in a ferret transmission model

Finally, we investigated the impact of the H18R and HA2-H111Y escape mutations and the putative permissive mutation, N133D, on viral replication and transmission in a ferret model (Fig. 7). Three donor ferrets were inoculated intranasally with 1×10^5^ 50% tissue culture infectious dose (TCID_50_) of each virus and three recipient ferrets were placed in adjacent cages the following day. Except one ferret inoculated with NL09-HA-H18R virus who was deceased at day six post-inoculation, all inoculated ferrets had mild clinical signs with normal body temperature (Supplementary Fig. 6). The peak viral titers of NL09-wt virus were 10-fold higher than NL09-HA-N133D virus in the inoculated ferrets, but this impact of N133D was not observed in the viruses harboring escape mutations (Fig. 7a). NL09-wt virus transmitted to two out of three recipient ferrets, with positivity first apparent at days 1 or 4 post-contact (Fig. 7b). NL09-HA-N133D transmitted to all recipients, with positivity detected uniformly at 1 day post contact (dpc) (Fig. 7c). Thus, the N133D mutation may enhance airborne transmissibility of NL09 virus. The viruses harboring the individual escape mutations H18R or HA2-H111Y spread to one out of three exposed ferrets, suggesting these mutations impose a fitness cost at the level of transmission (Fig. 7d,f). Likewise, viruses harboring the dual mutations, NL09-HA-H18R-N133D and NL09-HA2-H111Y-N133D, transmitted to one out of three exposed ferrets (Fig. 7e,g). Thus, while N133D reversed viral growth defects in human cell cultures, it is not sufficient to overcome the deleterious effects of the escape mutations for transmission in ferrets.

**Fig. 7.**
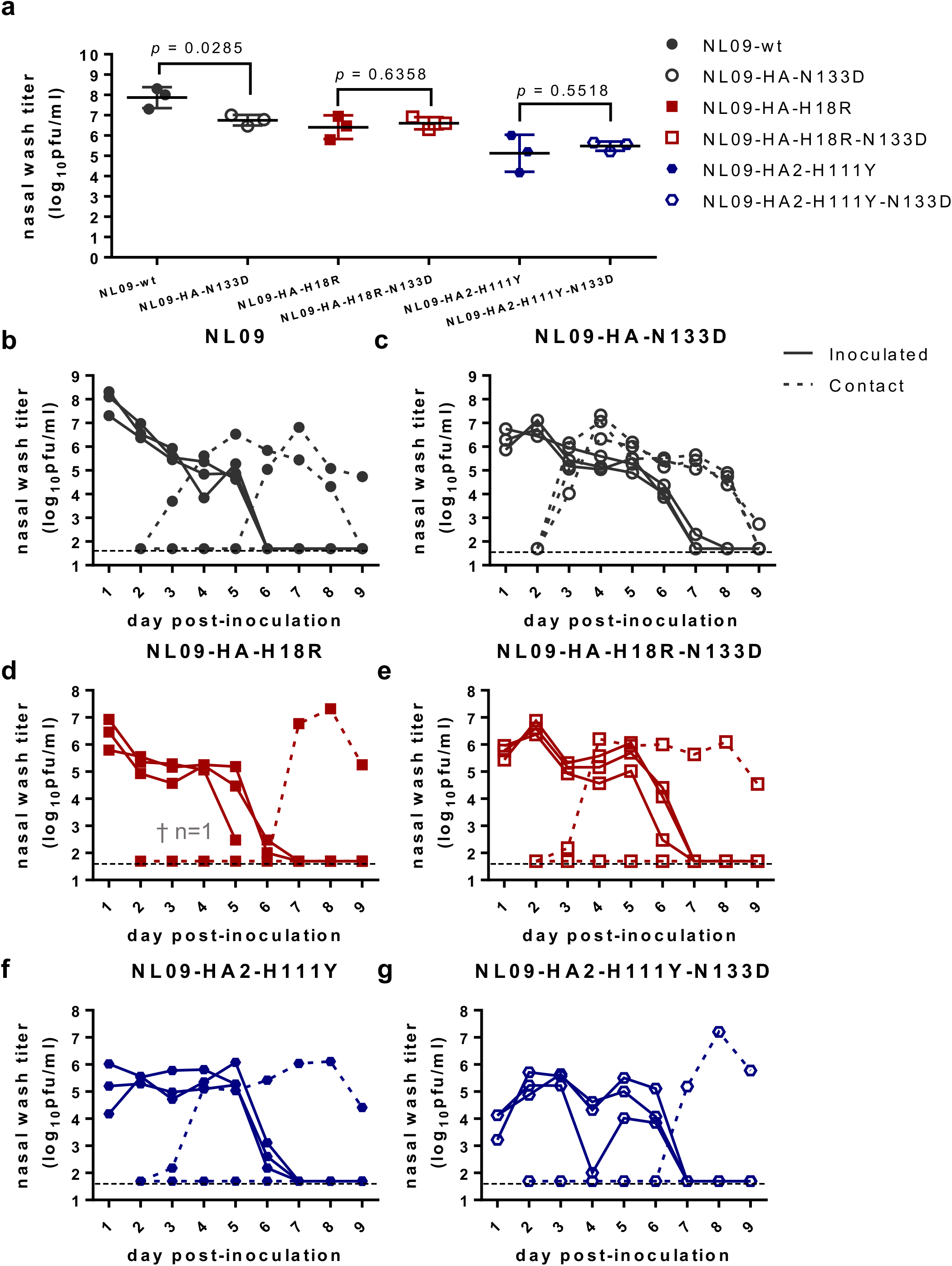
Ferret transmission of recombinant NL09 viruses harboring individual escape mutations with or without N133D. (a) Peak viral shedding titers observed in the inoculated ferrets. (b-g) Replication kinetics and airborne transmission of the indicated viruses. Statistical significance was evaluated using unpaired *t-test.* †, On day 6 post-inoculation, one inoculated ferret in the NL09-HA-H18R group was found unresponsive.

## Discussion

The HA stem domain shows high conservation among circulating IAVs, making it an attractive target for vaccination. This conservation is due at least in part to the importance of the stem region in the structure of the HA trimer and for HA-mediated membrane fusion. Here, we sought to probe the importance of functional constraints to conservation of the HA stem. We identified several escape mutations against two different stem-bnAbs, 70-1F02 and 05-2G02. All were located within the HA stem domain and abolished interaction with the antibodies; however, a range of fitness costs attributable to altered kinetics of fusion were apparent, suggesting an evolutionary trade-off between stem bnAb escape and efficient viral entry. While viral fitness in human cell cultures was restored by epistatic interaction with a mutation in the HA head domain, the permissive nature of this mutation was not apparent in a ferret transmission model. Taken together, the data point to a role for positive epistasis between the head and stem domains in overcoming fitness trade-offs, but underscore evolutionary constraint acting on the HA stem domain.

Evolutionary barriers to escape from immunological pressure can vary by protein location and in a strain specific manner. Antigenic drift in the HA stem is minimal compared to the head, consistent with reported differences in the mutational tolerance of these two domains^30, 31^. In addition, compared to H3 subtype IAV, H1 subtype viruses have a higher genetic barrier to evading stem binding antibodies^37^. Our data indicate that the H1 HA stem is capable of escape from immunological pressure in exchange for reduced fitness. While the observed negative fitness effects of stem variants may explain their rarity in nature, it is relevant to consider that the low fitness variants outcompeted the wild-type in the presence of the stem-bnAbs. These results are consistent with a recent human challenge study, in which a variant virus carrying an escape mutation was positively selected within an individual with pre-existing immunity^38^. Given individual-to-individual variation in immune status, IAVs are likely to experience fluctuations in selective pressure as they spread through a population. The potential for fixation of escape mutations in the HA stem is therefore likely to be reduced at a population level.

The escape mutations identified here show parallels with previously reported mutations that allow escape from other stem-bnAbs: H38R was observed during viral passage with a mouse monoclonal antibody, KB2^28^, while H111T or H111L allowed escape from C179 or CR6261 stem-bnAbs^15, 39^. Interestingly, we observed that S54P and H111Y in HA2, located in the interior of HA trimer, escape from both 70-1F02 and 05-2G02 stem-bnAbs. Likewise, A388T or V, which confer escape from several stem-bnAbs, is located in the interior of the HA trimer ^28, 38, 40^. Mutations within this interior region are likely to alter epitope recognition through indirect effects on HA conformation, which may explain the association with escape from multiple distinct antibodies. This mode of escape is notable in that it may allow viral evasion of polyclonal responses to the stem and distinct clonotypes present in different individuals.

We propose that positive epistasis facilitated the evolution of escape mutations in our experimental system. The deleterious side effects of antigenic variants, which are often stronger than the beneficial effects, constrain adaptation by preventing positive selection of antigenic variants. Fitness effects are context-dependent, however, such that an antigenic change may not incur costs if it occurs within the background of a permissive variant^41, 42, 43^. Comparing the frequency dynamics of variants in our passaged virus populations, we inferred that N133D arose ahead of the escape mutations. While N133D has negligible effects on viral fusion and stem-bnAb escape, it increases viral binding to host cell receptors. Notably, the addition of N133D to recombinant viruses harboring individual escape mutations enhanced replication in human cell cultures, yielding phenotypes comparable to that of NL09-wt virus. This pattern mirrors that seen in the evolution of oseltamivir resistance in the human seasonal H1N1 lineage in which permissive mutations in NA enabled the acquisition of resistance mutations that typically incurred fitness costs^44^. We therefore propose that the N133D mutation may act as a permissive mutation in the context of antibody escape. The recent sweep of HA N133D in pandemic H1N1 lineage IAVs circulating globally may therefore have important implications for further viral evolution.

Despite its beneficial effects in primary cell cultures, N133D did not enhance droplet transmission in ferrets. A recombinant virus harboring N133D transmitted rapidly to three out of three exposed animals, suggestive of similar or higher transmission fitness than NL09-wt virus. However, we did not see enhancement of airborne transmission when N133D was coupled to escape variants. This outcome was unexpected based on our observations in primary human cells and suggests that the permissive effects of N133D are insufficient to enable robust replication and transmission of escape variant viruses *in vivo.* The use of ferrets to model these processes presents a limitation, however. Although shown to be similar in prevalence of 2,3 and 2,6 linked sialylated glycans^45, 46, 47^, the distribution of specific glycan species likely differs between ferret and human respiratory tracts. These differences could lead to differing effects of N133D in the two hosts. Nevertheless, if mutations in the HA stem domain reduce transmissibility, as our ferret data suggest, they are unlikely to become prevalent in circulating viruses even if stem-targeting vaccines introduce increased immune pressure within the population. However, our study is not exhaustive and was not designed to select for mutations that act at the level of transmission. Mutations that ameliorate the transmission fitness effects of stem variants may emerge under relevant selective conditions in humans.

In summary, our data suggest that evolution of circulating IAV in response to increased selective pressure on the HA stem domain would likely be highly constrained. However, epistasis may increase the potential for emergence of variants that escape neutralization by stem-bnAbs. Therefore, active surveillance to monitor viral antigenic evolution will be needed along with universal vaccine development.

## Acknowledgements

The research was funded by the NIH/NIAID Centers of Excellence in Influenza Research and Surveillance (CEIRS), contract number HHSN272201400004C and R01 AI165644 to ACL.

## Author contributions

C-YL contributed to the conception of work, experimental design, data acquisition and analysis, interpretation of data and led preparation of the manuscript; CJC, GG, BS, FCF, LCG, LMF and DK contributed to data acquisition; JW provided critical reagents and expertise; GST contributed to experimental design and data and funding acquisition; DRP contributed to conception of the work and experimental design; ACL contributed to conception of work, funding acquisition, experimental design and data analysis and interpretation. All authors contributed to the writing of the manuscript.

## Methods

### Cells, viruses, and antibodies

Madin-Darby canine kidney (MDCK) cells were maintained in minimal essential medium (MEM) (Gibco, USA) supplemented with 10% fetal bovine serum (FBS) (R&D SYSTEMS, USA) and penicillin-streptomycin (PS) (Gibco, USA). Baby Hamster Kidney (BHK-21) cells were maintained in MEM with GlutaMAX™ Supplement (Gibco, USA), 5% FBS and PS. 293T cells were maintained in Dulbecco’s Modified Eagle Medium (DMEM) (Gibco, USA) supplemented with 10% FBS and PS. Normal human bronchial epithelial (NHBE) cells (Lonza, Switzerland) were amplified and differentiated into air–liquid interface cultures as recommended by Lonza and described previously^49^. Influenza A/Netherlands/602/2009 (H1N1) [NL09] virus was generated using reverse genetics. Briefly, 293T cells transfected with reverse genetics plasmids 24 hours previously were co-cultured with MDCK cells at 37 °C for 48 hours. Recovered virus was plaque purified and propagated in MDCK cells to generate a working stock. 70-1F02 and 1009-3E04 antibodies were isolated from patients who were naturally infected with 2009 pandemic H1N1 virus and 05-2G02 antibody was isolated from an immunized individual ^20 24^.

### Selection of escape mutants

To isolate escape variants, NL09 virus was serially passaged 10 times with a range of the stem bnAbs. First, single replicates with an initial concentration of 20, 2, or 1 μg/mL of 70-1F02 or 05-2G02 were mixed with an MOI=1 PFU/cell of NL09 virus for one hour at 37 °C. Next, the mixtures were inoculated onto a 6-well plate of MDCK cells and incubated for 2 days at 37 °C. The supernatant was collected, mixed with increased concentration of the stem-bnAbs, and inoculated onto fresh MDCK cells. During serial passage with the stem-bnAbs, the antibody concentration was increased 2-fold if the infected cells showed gross cytopathic effects (CPE) (70-90% floating cells) ^32^. If CPE was moderate to mild, the antibody concentration was maintained. As a control, viruses without stem-bnAbs were passaged in parallel in MDCK cells. The populations passaged 10 times and the early passaged populations which showed moderate resistance to stem-bnAb were deep-sequenced to monitor variant dynamics. In addition, six plaques were picked from the passaged populations and sequenced to verify the deep sequencing results and to evaluate co-occurrence of mutations (Supplementary Table 1).

### Next generation sequencing and variant analysis

Analysis of non-consensus variants was made using LoFreq^50^ following the Genome Analysis Toolkit best practices^51^. In brief, after removing adapters using Trimmomatic (version 0.39)^52^, reads were aligned to their reference sequence using the option mem from BWA^53^. Data formatting for GATK was made using Picard (http://broadinstitute.github.io/picard/). Reads were realigned using RealignerTargetCreator and IndelRealigner from GATK. The quality of bases was recalculated using BaseRecalibrator from GATK. The resulting bam file was used to perform variant calling analysis by LoFreq. Only variants at a frequency of 0.05 with a coverage equal or above 400 were used.

### Site-directed mutagenesis and recombinant virus rescue

All viruses used in this study were generated by reverse genetics^54^. 293T cells were transfected with eight bi-directional plasmids encoding PB2, PB1, PA, HA, NA, NP, M, and NS genes. After 24 hours post-transfection, 1 ×10^6^ cells of MDCK cells were overlaid with virus media containing 1 μg/mL TPCK treated Trypsin. Rescued viruses were plaque purified and passaged once in MDCK cells. The resultant cell passage 1 stocks were used in experiments. Single amino acid mutations were introduced into the NL09 HA by site-directed mutagenesis using primers (Supplementary Table 2). Unique biological materials such as recombinant viruses are available from the corresponding author upon request.

### Syncytia formation assay

BHK-21 cells were transfected with 1 μg of pCAGGS plasmid encoding the NL09 wild-type or escape mutant HA proteins using Lipofectamine (Invitrogen, USA) and Plus reagent (Invitrogen, USA). At 24 h post-transfection, HA-expressing cells were washed once with PBS and treated with 2.5 μg/mL of TPCK-treated trypsin (Sigma-Aldrich, USA) for 10 minutes. Afterward, the cells were exposed to pH adjusted PBS for 5 minutes at 37 °C. The cells were neutralized by complete growth media (MEM, supplemented with GlutaMAX™ Supplement, 5% FBS and PS) and incubated for two hours at 37 °C to allow syncytia formation. The cells were fixed and stained with the Hema3Stat Pak (Fisher Scientific, USA) according to the manufacturer’s protocol. Syncytia were visualized and photographed using a Zeiss Axio Observer inverted microscope with an attached digital camera (Zeiss, Germany).

### Cell-based ELISA

BHK-21 cells in 96-well plate were transfected with 100 ng of pCAGGS NL09 HA (wild-type or escape mutants) using Lipofectamine 3000 (Invitrogen, USA). At 24 h post-transfection, the HA-expressing cells were washed twice with PBS and fixed with 4% formaldehyde for two hours. The fixed cells were washed twice with PBS and blocked with 5% skim milk in PBS with 0.05% Tween20 for one hour. Serially diluted stem-bnAbs were added on to the plate and incubated for one hour at room temperature, and then washed 4 times with PBST (PBS+0.05% Tween20). The secondary antibody (anti-human IgG (H+L)-HRP conjugated, Promega, USA) was added to the plate and incubated for one hour. After washing with PBST four times, the plates were treated with 100 μL of TMB solution (Thermo Fish Scientific, USA) and incubated for 15 minutes. The reactions were halted by addition of 2 N sulfuric acid. The color was read at 450 nm with the reference wavelength of 655 nm.

### Viral fusion kinetics

MDCK cells in a 96-well plate were infected at an MOI=1 PFU/cell. At 24 h post-infection, the cells were washed twice with PBS and treated with 5 μg/mL TPCK-treated trypsin for 10 min at 37 °C. The trypsin was neutralized by complete growth media (MEM supplemented with 10% FBS and PS). After washing with PBS, the cells were incubated with 1 μg/mL of 1009-3E04 antibody for one hour at 37 °C. Following the antibody treatment, the cells were washed twice with PBS containing 2% FBS (PBSF) and then treated with pH adjusted PBS at pH=5.0 for 0, 2, 4, 6, 8, and 10 minutes at 37 °C. Following the acid treatment, the cells were washed with PBSF, then incubated with anti-human IgG-HRP for one hour at room temperature. After washing with PBSF, the cells were treated with 100 μL of TMB solution and incubated for 15 minutes. The reactions were halted by addition of 2 N sulfuric acid. The color was read at 450 nm with the reference wavelength of 655 nm.

### Multi-step viral growth kinetics in MDCK, A549, and NHBE cells

MDCK and A549 cells were seeded onto 6-well plate. After 24 h incubation, the cells were washed three times with PBS and inoculated with MOI 0.002 PFU/cell of NL09 wild-type or NL09 escape mutant viruses for MDCK cells and MOI 0.01 PFU/cell for A549 cells. After 1 h absorption at 4 °C, the cells were washed three times with PBS and virus media with 1 μg/ml TPCK treated trypsin (Sigma-Aldrich, USA) was added. A 200 μL volume of supernatant was harvested at each time-point with replacement of the same volume of virus medium. NHBE cells were differentiated at an air-liquid interface. After the mucus was removed from the apical surface, the cells were washed five times with PBS and inoculated with MOI 0.01 of NL09 wild-type or NL09 escape mutant viruses.

### Solid phase binding assay

Serially diluted (20, 10, 5, 2.5, 1.25, 0.625, and 0.3125 μg/mL) biotinylated glycan 3’SLN-PAA and 6’SLN-PAA (Carbosynth, USA) in PBS were added to the wells of streptavidin-coated high binding capacity 96-well plates (Thermo Fisher Scientific, USA). After incubation overnight at 4 °C, the plates were washed with PBST and blocked with PBS containing 2% BSA for two hours at room temperature, then washed with PBST. 10^6^ PFU of virus was prepared in PBS containing 10 μM GS4071 neuraminidase inhibitor and added to the wells and incubated overnight at 4 °C. The plates were washed with PBST and incubated with 5 μg/mL 1009-3E04 monoclonal antibody for an hour. After washing with PBST, the wells were incubated with secondary antibody (anti-human IgG (H+L)-HRP conjugated, Promega, USA) for an hour, then washed with PBST again. The plates were treated with 100 μL of TMB solution and incubated for 15 minutes. The reactions were halted by addition of 2 N sulfuric acid. The color was read at 450 nm with the reference wavelength of 655 nm.

### Glycan microarray

The viruses were purified through a 25% sucrose cushion, then labeled with Alexa Fluor 488 (Invitrogen, USA) as previously described^55, 56^. Labeled viruses were dialyzed against PBS using a 7,000-molecular-weight-cutoff Slide-A-Lyzer mini-dialysis unit (Thermo Fisher Scientific, USA) overnight at 4 °C. The labeled viruses were run on v4.0 of the CFG Glycan Array using the buffers and conditions described previously^55, 56^.

### Ferret transmission experiment

Ferret studies were approved and conducted in compliance with all the regulations stated by the Institutional Animal Care and Use Committee (IACUC) of the University of Georgia (AUP2019 03-021-Y3-A7). Studies were conducted under BSL-2 conditions at the Poultry Diagnostic and Research Center (PDRC), University of Georgia. Animal studies and procedures were performed according to the Institutional Animal Care and Use Committee Guidebook of the Office of Laboratory Animal Welfare and PHS Policy on Humane Care and Use of Laboratory Animals. Animal studies were carried out in compliance with the ARRIVE guidelines (https://arriveguidelines.org). Twenty-week-old female ferrets were acquired from Triple F Farms (Gillett, PA, USA). The next day after arrival, ferrets were anesthetized with an i.m. injection (0.5 ml/kg) of a ketamine cocktail (20 mg/kg Ketamine, 1 mg/kg Xylazine) in the thighs, followed by blood collection to test serum samples for prior IAV infection using NP ELISA (IDEXX, ME, USA). All ferrets tested negative for prior IAV exposure. A subcutaneous implantable temperature transponder (BMDS, DE, USA) was inserted in the back of the neck to ID and monitor body temperature of each ferret. Ferrets were acclimated for 7 days upon arrival before virus challenge. For each virus evaluated, ferrets (n=3) were anesthetized as described above and inoculated with 1×10^5^ TCID_50_/ferret in a final volume of 1 ml in 1X PBS. On day 1 post-inoculation (dpi), the direct inoculated ferrets were placed with naïve respiratory contact ferrets (ratio 1:1/isolator). Temperature, weight, and clinical signs were monitored every day. Nasal wash samples were obtained for all the ferrets every day until 9 dpi (8 days post-contact, dpc) by introducing 1 ml volume of 1X PBS-BSA 1% into the nasal passages to induce sneezing, collecting the aspirate using a sterile plastic dish, and washing the plate with an additional 1 ml of 1X PBS supplemented with antibiotic/antimycotic solution (Gibco, USA). Nasal washes were then aliquoted and frozen at −80°C until further processing. To avoid direct-contact contamination, 70% EtOH surface decontamination of gloves and equipment was performed between handling inoculated and contact animals. Contact animals were always handled before inoculated animals. Different bite-resistant gloves, utensils, and tools were used between contact and virus-inoculated animals, with disposable gowns and gloves changed between isolators. On 14 dpi, all ferrets were anesthetized as described above, bled, and humanely euthanized using 1 ml of Euthasol^®^ (Virbac, TX, USA).

### Sequence analysis

3341 HA sequences of 2009 pandemic H1N1 virus isolates from 2009 to 2021 were collected in GISAID (Global Initiative on Sharing All Influenza Data) on September 25^th^, 2021 and sorted by years in Excel.

### Data availability

Raw data used for the generation of Figures 1-7 and Supplementary Figures 1-6 will be available before publication. All the other data that support the conclusions of the study are available from the corresponding author upon request.

