## Supplemental Information for "Epistasis reduces fitness costs of influenza A virus escape from stem-binding antibodies"

### Supplementary Table 1. Mutations found by plaque purification of passaged virus populatio n

| a.a[a] | H3 [b] numbering | NL09 (WT) | 07-1F02 | | | | | | 50-2G02 | | | | | | No bnAb | | | |
|---|---|---|---|---|---|---|---|---|---|---|---|---|---|---|---|---|---|---|
| | | | Rep 1 | | Rep 2 | | Rep 3 | | Rep 1 | | Rep 2 | | Rep 3 | | Rep 1 | Rep 2 | Rep 3 | Rep 4 |
| | | | Early (P3) | P10 | Early (P7) | P10 | Early (P7) | P10 | Early (P3) | P10 | Early (P4) | P10 | Early (P6) | P10 | P10 | P10 | P10 | P10 |
| 16 | - | N | | | | | | | | | | | | | | **T (2/4)** | | |
| 25 | 18 | H | | **R (6/6)**[c] | | | | | | | | | | | | | | |
| 28 | 21 | N | | | | | | | | | | **N (6/6)** | | | | | | |
| 44 | 37 | T | | | | | | | | | | | | | | | | |
| 45 | 38 | H | | | **R (1/6)** | **R (5/6)** | | | | | | | | | | | | |
| 136 | - | K | | | | | | | | | | | | | | | | |
| 141 | 128 | P | | | | | | | | | | | | | | | | |
| 146 | 133 | N | | **D (6/6)** | **D (4/6)** | **D (6/6)** | **D (6/6)** | **D (6/6)** | | | **D (2/5)** | **D (4/4)** | **D (3/6)** | **D (5/5)** | **D (1/6)** | | | |
| 152 | 138 | A | | | | | | | | | | | | | | | | |
| 158 | 144 | A | | | | | | | | | **V (1/5)** | | | | | | | |
| 172 | 158 | G | | | | | | | | | | | | | | | **E (2/2)** | |
| 173 | 159 | N | | | **D (4/6)** | | | | | | | | **D (3/6)** | | | | | **D (2/3)** |
| 199 | 185 | P | | | | | | | | | | | | | | | | |
| 207 | 193 | S | | | | | | | | | | | | | | | | |
| 212 | 198 | A | | | | | | | | | | | | | | **E (1/4)** | | |
| 223 | 209 | Y | | | | | | | | | | | | | | **H (1/4)** | | |
| 253 | 239 | P | | | | | | | **Q (1/4)** | | | | | | | | | |
| 287 | 272 | T | | | | | | | | | | **A (6/6)** | | | | | | |
| 333 | 318 | T | | | | | | | | **K (2/6)** | | | | | | | | |
| 335 | 320 | L | | | | **W (1/6)** | | | | | | | | | | | | |
| 381 | 37 (HA2) | D | | | **G (4/6)** | | | | | | | | | | | | | |
| 388 | 44 | A | **V (6/6)** | | **T (1/6)** | | | | **V (4/4)** | | | | | | | | | |
| 398 | 54 | S | | | | | | **P (6/6)** | | | | | | | | | | |
| 447 | 103 | E | | | | | | | | | | | **K (1/6)** | | | | | |
| 455 | 111 | H | | | | | | | | **Y (4/6)** | | | | **Y (5/5)** | | | | |
| 458 | 114 | S | | | | | | | | | | | | | | | | |
| 493 | 149 | M | | | | | | | | | | | | | | | | |
| 515 | 171 | E | | | **K (1/5)** | **K (5/6)** | | | | | | | | | | | | |
| 544 | 300 | V | | | | | | | | | | | | | | | | |

[a] Amino acid numbering from start codon
[b] H3 numbering
[c] Frequency of mutation among sequenced plaque purified viruses (no. with mutations present / no. plaques analyzed).

**Supplementary Table 2. STAR Methods oligonucleotide list, primers used for site-directed mutagenesis**

| Oligonucleotides | | |
|---|---|---|
| QC_NL09_HA_H25R_F | gacacattatgtataggttatcgtgcgaacaattcaacagacact | N/A |
| QC_NL09_HA_H25R_R | agtgtctgttgaattgttcgcacgataacctatacataatgtgtc | N/A |
| QC_NL09_HA_N28T_F | gtataggttatcatgcgaacacttcaacagacactgtagacac | N/A |
| QC_NL09_HA_N28T_R | gtgtctacagtgtctgttgaagtgttcgcatgataacctatac | N/A |
| QC_NL09_HA_H45R_F | gaaaagaatgtaacagtaacacgctctgttaaccttctagaagac | N/A |
| QC_NL09_HA_H45R_R | gtcttctagaaggttaacagagcgtgttactgttacattcttttc | N/A |
| QC_NL09_HA_T287A_F | ggatctggtattatcatttcagatgcaccagtccacga | N/A |
| QC_NL09_HA_T287A_R | tcgtggactggtgcatctgaaatgataataccagatcc | N/A |
| QC_NL09_HA_T333K_F | gcacaaaattgagactggccaaaggattgaggaatgtc | N/A |
| QC_NL09_HA_T333K_R | gacattcctcaatcctttggccagtctcaattttgtgc | N/A |
| QC_NL09_HA_L335W_F | gactggccacaggatggaggaatgtcccg | N/A |
| QC_NL09_HA_L335W_R | cgggacattcctccatcctgtggccagtc | N/A |
| QC_NL09_HA_S398P_F | ccattgacgagattactaacaaagtaaatcctgttattgaaaagatgaataca | N/A |
| QC_NL09_HA_S398P_R | tgtattcatcttttcaataacaggatttactttgttagtaatctcgtcaatgg | N/A |
| QC_NL09_HA_H455Y_F | gaaaatgaaagaactttggactactacgattcaaatgtgaagaacttat | N/A |
| QC_NL09_HA_H455Y_R | ataagttcttcacatttgaatcgtagtagtccaaagttctttcattttc | N/A |
| QC_NL09_HA_E515K_F | cagaggaagcaaaattaaacagaaaagaaatagatggggtaaagctg | N/A |
| QC_NL09_HA_E515K_R | cagctttaccccatctatttcttttctgtttaattttgcttcctctg | N/A |
| QC_NL09_D381G_F | caggatatgcagccggcctgaagagcacaca | N/A |
| QC_NL09_D381G_R | caggatatgcagccggcctgaagagcacaca | N/A |
| QC_NL09_A388V_F | caggatatgcagccggcctgaagagcacaca | N/A |
| QC_NL09_A388V_R | caggatatgcagccggcctgaagagcacaca | N/A |
| QC_NL09_A388T_F | caggatatgcagccggcctgaagagcacaca | N/A |
| QC_NL09_A388T_R | caggatatgcagccggcctgaagagcacaca | N/A |
| QC_NL09_E447K_F | caggatatgcagccggcctgaagagcacaca | N/A |
| QC_NL09_E447K_R | caggatatgcagccggcctgaagagcacaca | N/A |
| QC_NL09_A158V_F | caggatatgcagccggcctgaagagcacaca | N/A |
| QC_NL09_A158V_R | caggatatgcagccggcctgaagagcacaca | N/A |
| QC_NL09_P253Q_F | caggatatgcagccggcctgaagagcacaca | N/A |
| QC_NL09_P253Q_R | caggatatgcagccggcctgaagagcacaca | N/A |

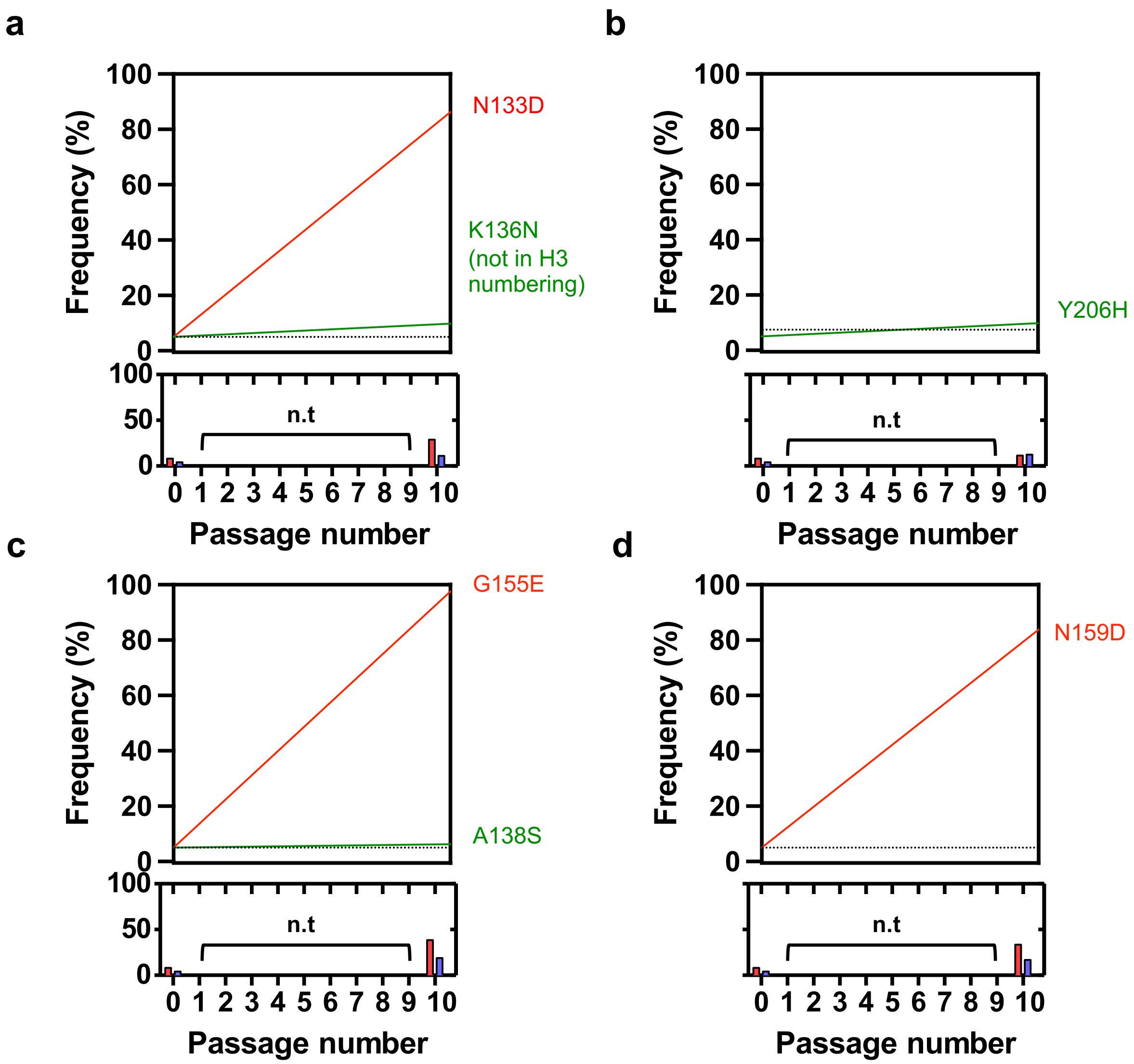


**Supplementary Fig. 1. Dynamics of mutation frequency and resistance of NL09 virus popula tions passaged without antibodies. (a-d)** MOI=1 PFU/cell of NL09 virus were initially inoculated onto MDCK cells, then the supernatants were blindly passaged 10 times in MDCK cells. n.t = not tested; dotted line = limit of detection (5%).

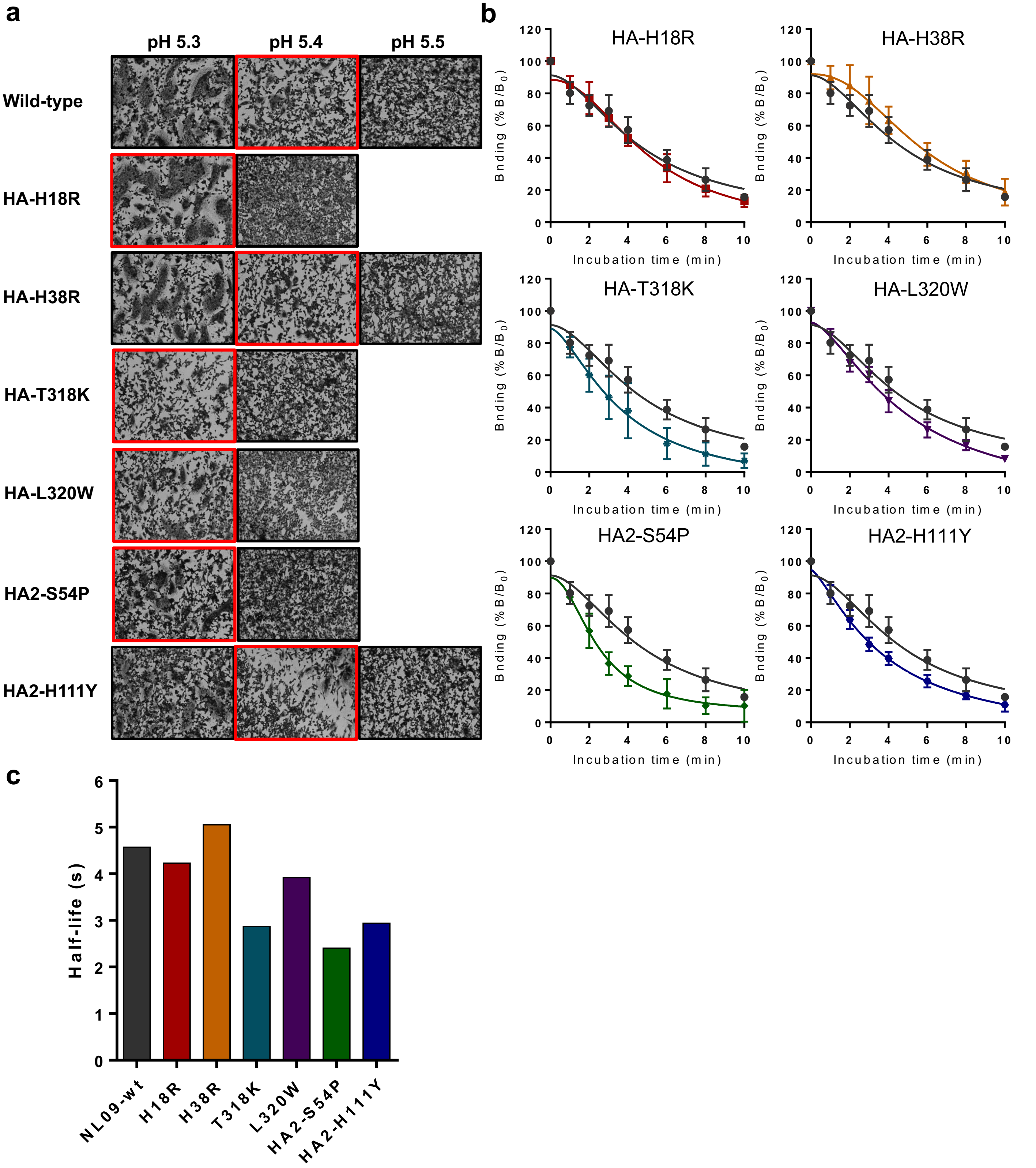


**Supplementary Fig. 2. Effect of escape mutations on virus and membrane fusion.** (a) Fusion pH of NL09 HA harboring individual escape mutations was measured by syncytium formation. Red rectangles mark the highest pH at which syncytia were observed. (b) Kinetics of low pH-induced co nformational change of NL09 HA proteins containing single escape mutations, as measured by ELI SA with a conformation-sensitive antibody. (c) Time to 50% dissociation of the conformation-sensiti ve antibody, derived from curves in B using nonlinear regression analysis.

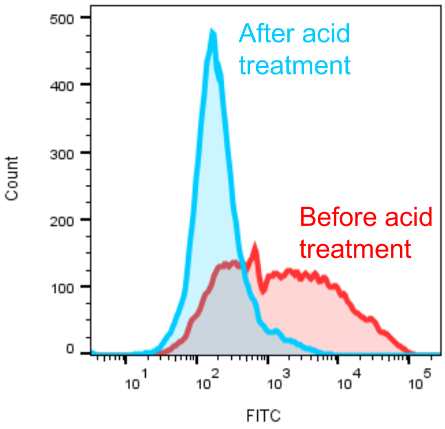


**Supplementary Fig. 3. Antibody detachment by pH induced HA conformational change.** MOI=1 PFU/cell of NL09-wt virus was inoculated onto MDCK cells, and the cells were trypsinized a nd collected at 24 h post-inoculation. The cells were incubated with 1009-3E04 antibody for one ho ur, then they were divided into two groups; one treated with pH=5.0 of acidic PBS for 10 minutes, a nd one that remained untreated. Antibody binding was analyzed using flow cytometry.

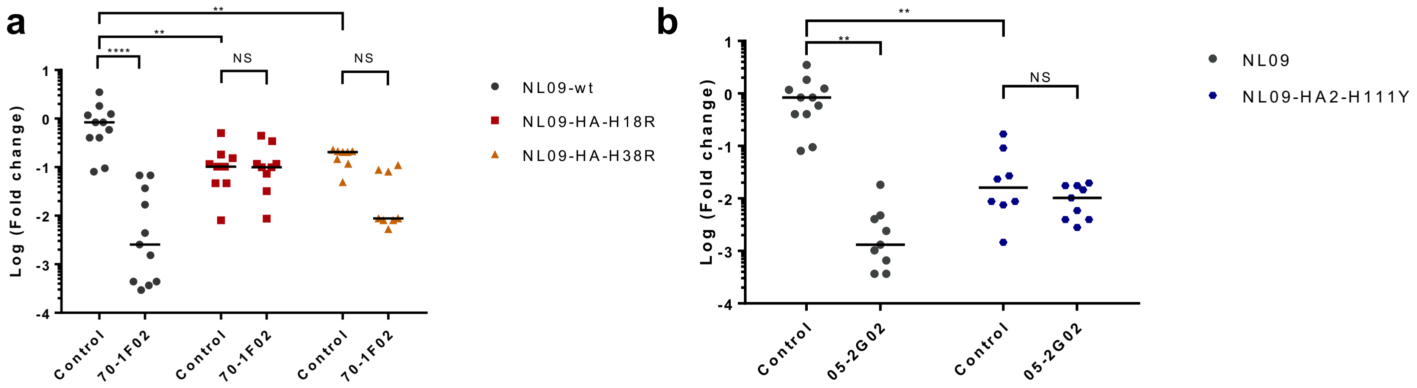


**Supplementary Fig. 4. Growth inhibition by stem-bnAbs was relieved by individual escape mutations.** Viral yield at 48 h post-inoculation in NHBE cells is shown in the absence or presence of stem-bnAbs (a) 70-1F02 or (b) 05-2G02. Each data point corresponds to an independent experi ment. Statistical significance was assessed using a one-way ANOVA with Tukey's multiple comparisons test. ****, < 0.0001; ** < 0.01; NS, Not significant.

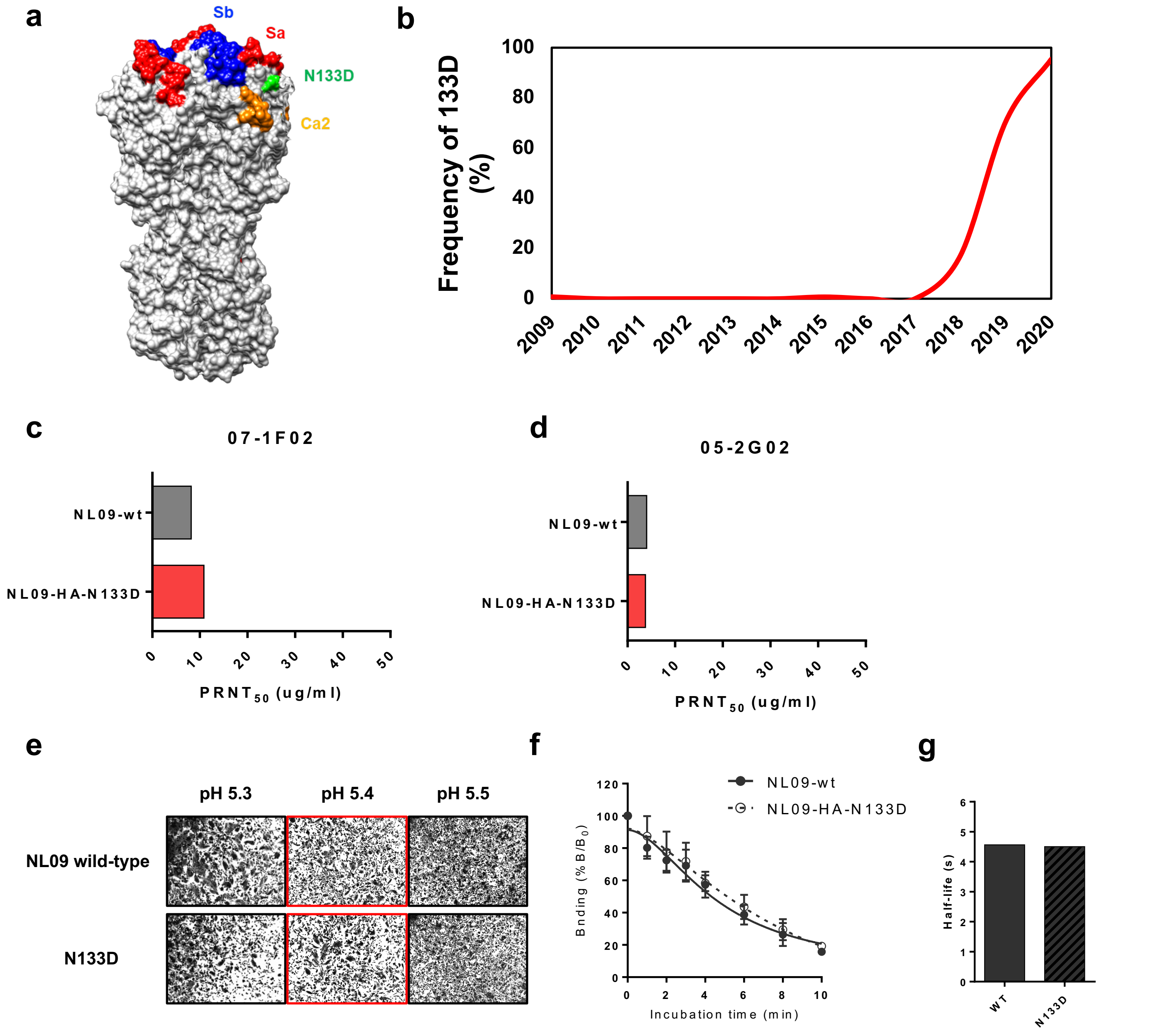


**Supplementary Fig. 5. N133D mutation does not affect resistance to stem-bnAbs nor HA fus ion activity.**

(a) Location of residue 133 on the HA of A/California/04/2009(H1N1) (PDB ID 3LZG) (b) Occurrenc e and dominance of HA 133D among circulating 2009 pandemic H1N1 strains. 3341 HA sequences of 2009 pandemic H1N1 virus isolates were downloaded from GISAID (Global Initiative on Sharing All Influenza Data) on September 25th, 2021 and sorted by years. (c and d) Resistance of NL09-HA -N133D toward the two stem-bnAbs were measured using plaque reduction neutralization test. (e) Fusion pH of NL09-HA-N133D were measured using syncytia formation assay. (f) Fusion kinetics o f NL09-HA-N133D were determined using ELISA. (g) Time to 50% dissociation of the antibody. The half-life of the fusion kinetics of NL09-HA-N133D was calculated using nonlinear regression analysi s.

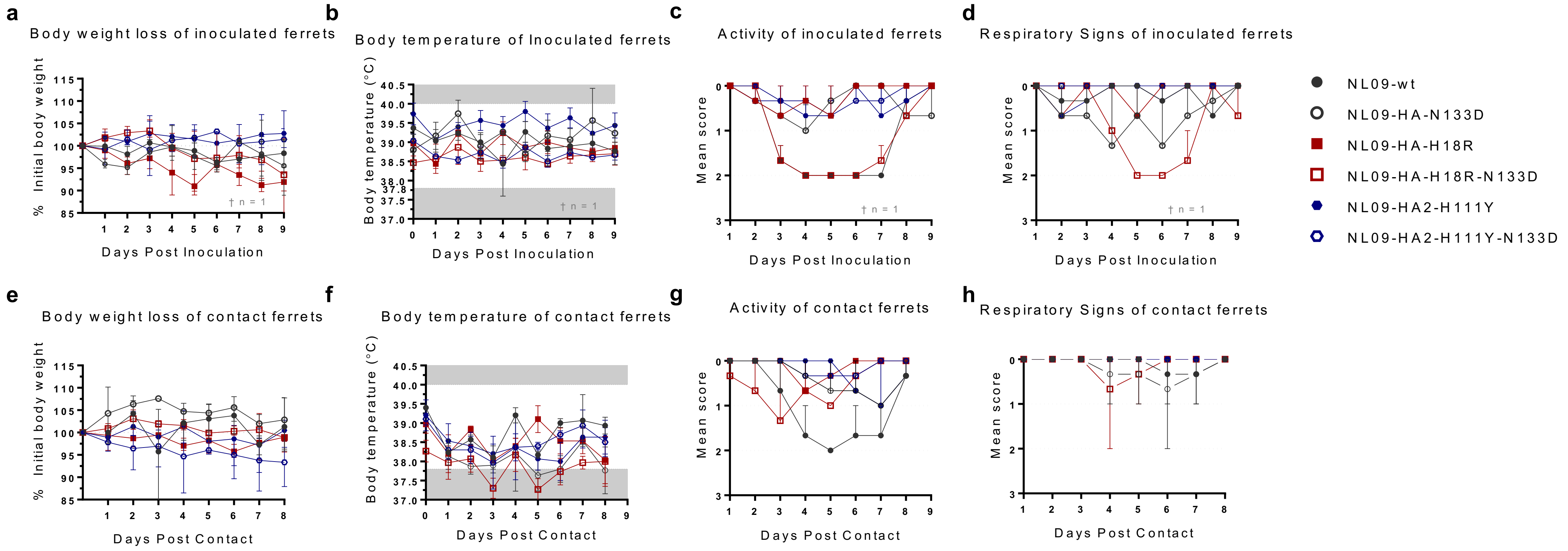


**Supplementary Figure 6. Clinical signs associated with mutant H1N1 pdm influenza A viruses in ferrets.** Ferrets were inoculated intranasally ( n=3/group, panels a-d) with $10^5$ $TCID_{50}$/animal of either NL09-wt virus or one of the recombinant viruses listed above. Inoculated ferrets were housed in HEPA-filtered isolators (n=1/isolator, 3 isolators/group). Respiratory contact ferrets ( n=3/group, at 1:1 ratio Inoculated:Contact, panels e-h) were introduced at 1 dpi. Ferrets were monitored daily for body weight changes (a and e), body temperature changes (b and f), activity score (c and g; 0=normal, 1=mild, 2=moderate, 3=no activity), respiratory signs ( d and h; 0=normal, 1=mild, 2=moderate, 3=severe). Normal body temperate for ferrets falls within 37.8ºC and 40ºC. †: On day 6 post-inoculation, one inoculated ferret in the NL09-HA-H18R group was found unresponsive: Post-mortem findings included presence of abundant catarrhal exudate in the nasal passages, laryngeal isthmus, and larynx compatible with an upper respiratory infection. Small amount of exudate was also observed in the left tympanic bulla and is suggestive of otitis.
